# BMAL1-HIF2α heterodimers contribute to ccRCC

**DOI:** 10.1101/2024.06.07.597806

**Authors:** Rebecca M. Mello, Diego Gomez Ceballos, Colby R. Sandate, Daniel Agudelo, Celine Jouffe, Nina Henriette Uhlenhaut, Nicolas H. Thomä, M. Celeste Simon, Katja A. Lamia

## Abstract

Circadian disruption enhances cancer risk, and many tumors exhibit disordered circadian gene expression. We show rhythmic gene expression is unexpectedly robust in clear cell renal cell carcinoma (ccRCC). Furthermore, the clock gene *BMAL1* is higher in ccRCC than in healthy kidneys, unlike in other tumor types. BMAL1 is closely related to ARNT, and we show that BMAL1-HIF2α regulates a subset of HIF2α target genes in ccRCC cells. Depletion of *BMAL1* reprograms HIF2α chromatin association and target gene expression and reduces ccRCC growth in culture and in xenografts. Analysis of pre-existing data reveals higher *BMAL1* in patient-derived xenografts that are sensitive to growth suppression by a HIF2α antagonist (PT2399). We show that BMAL1-HIF2α is more sensitive than ARNT-HIF2α to suppression by PT2399, and increasing *BMAL1* sensitizes 786O cells to growth inhibition by PT2399. Together, these findings indicate that an alternate HIF2α heterodimer containing the circadian partner BMAL1 contributes to HIF2α activity, growth, and sensitivity to HIF2α antagonist drugs in ccRCC cells.

## Main

Many tumors exhibit disruption of circadian rhythms ^1^, and deletion of the clock component BMAL1 exacerbates tumor burden in several genetically engineered mouse models of cancer ^2,3^. However, circadian disruption is not universally observed in cancer cells, and BMAL1 depletion improves outcomes in some cancer models ^4^. It has been unclear why genetic deletion of BMAL1 enhances the growth of some tumors and suppresses others.

The von Hippel Lindau (VHL) ubiquitin ligase is inactivated in 50-85% of clear cell renal cell carcinomas (ccRCC) ^5–8^. VHL targets hypoxia inducible factors 1 alpha (HIF1α) and 2 alpha (HIF2α, a.k.a. EPAS1) for degradation ^9^. HIF1α and HIF2α are basic helix-loop-helix and PER-ARNT-SIM domain (bHLH-PAS) transcription factors that bind DNA with a common heterodimer partner HIF1β (a.k.a. ARNT), and increase the expression of genes involved in metabolism, proliferation, and angiogenesis ^7,8,10–12^. Suppression of HIF2α is required for VHL to inhibit ccRCC tumor growth ^13,14^, highlighting the oncogenic role of HIF2α in ccRCC.

In mammals, circadian clocks comprise a transcription-translation feedback loop, centered around the heterodimeric transcription factor complex containing CLOCK and BMAL1 ^15^. CLOCK and BMAL1 are bHLH-PAS transcription factors and are closely related to ARNT and HIF2α ^16,17^ (Fig. 1A,B). At the time of its initial characterization, BMAL1 was found to be dispensable for developmental processes in which HIFs are key players, and was therefore considered not to be a relevant partner for HIF alpha subunits ^18–21^. This impression was reinforced when X-ray crystal structures described divergent arrangements of the bHLH and PAS domain interfaces for CLOCK-BMAL1 and for HIFα-ARNT complexes ^16^. However, the arrangement of BMAL1 PAS domains is flexible even within CLOCK-BMAL1 heterodimers ^22^, and BMAL1 can activate transcription via hypoxia response elements (HREs) in cooperation with HIF alpha subunits ^23,24^. Accumulating evidence indicates that BMAL1 is an important partner for HIF1α-dependent hypoxic responses ^23–25^. Together, these findings motivate a reconsideration of the possible physiological relevance of a more diverse set of bHLH-PAS heterodimer pairings.

**Figure. 1.**
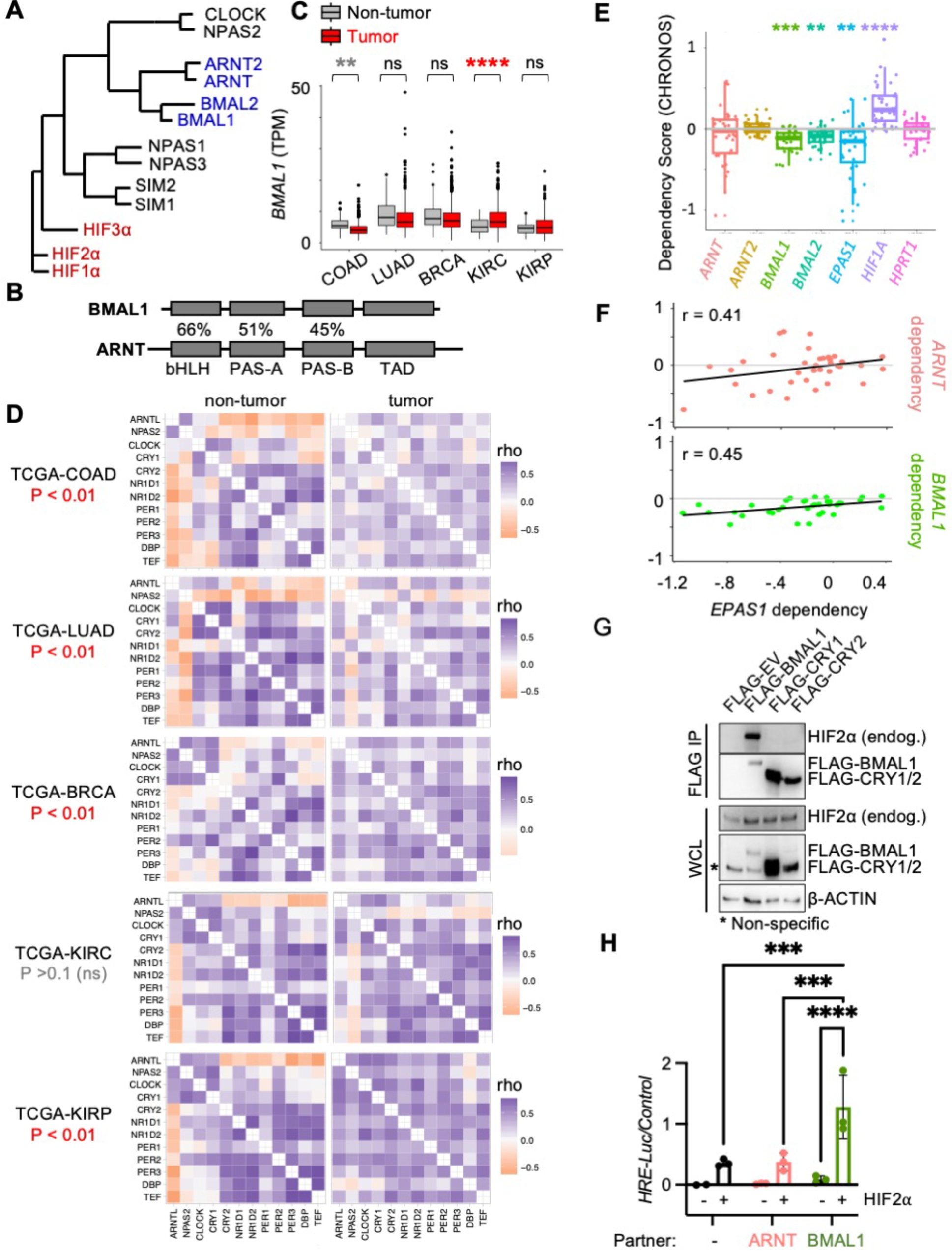
BMAL1 forms an active heterodimer with HIF2α. (**A**) Phylogenetic tree for bHLH-PAS proteins. (**B**) Percent sequence identity for bHLH and PAS domains in BMAL1 and ARNT. (**C,D**) Detection of *BMAL1* (transcripts per million, TPM) (*C*) and clock correlation distance (CCD) heatmaps (*D*) calculated from RNA sequencing data from tumors and adjacent normal tissues in cancer genome atlas projects: colorectal adenocarcinoma (COAD), lung adenocarcinoma (LUAD), breast cancer (BRCA), kidney clear cell renal cell carcinoma (KIRC), and renal papillary cancinoma (KIRP). (**E,F**) Dependency (CHRONOS) scores (*E*) and correlations thereof (*F*) for bHLH-PAS members in RCC cell lines from DepMap ^29,30^. (**G**) Detection of indicated proteins in whole cell extracts (input) or following immunoprecipitation of the FLAG tag from 786O cells transiently expressing the indicated plasmids. (**H**) Relative luciferase units detected from U2OS cells expressing *HRE-Luciferase* and additional indicated plasmids with (red) or without (black) exogenous stabilized HIF2α (P405A, P531A, N837A). In (*C,E*) boxplots depict the median and interquartile range (IQR), whiskers extend either to the minimum or maximum data point or 1.5*IQR beyond the box, whichever is shorter. Outliers (values beyond the whisker) are shown as dots in (*C*). In (*H*) bars represent mean ± s.e.m. for three experimental replicates and symbols represent the mean of n=5 measurements for each experiment. In (*C,E,H*) ** Padj < 0.01, *** P< 0.001, **** P < 0.0001 by two-way ANOVA with Tukey’s (*C*,*E*) or Sidak’s (*H*) correction for multiple comparison.

Small molecules that interact with a pocket in the PAS-B domain of HIF2α disrupt the formation of HIF2α heterodimers and are used to treat ccRCC. Variability in responses to these drugs can be caused by mutations surrounding their binding site in HIF2α or ARNT in some cases but is not generally understood ^14,26–28^. Here, we demonstrate that BMAL1 forms a transcriptionally active heterodimer with HIF2α in ccRCC-derived cells and contributes to HIF2α-driven gene expression, cell and tumor growth, and sensitivity to growth suppression by the HIF2α antagonist PT2399.

### ccRCC tumors maintain robust circadian rhythms

Using data from the Clinical Proteomic Tumor Analysis Consortium (CPTAC), and the Cancer Genome Atlas (TCGA), we find that *BMAL1* expression is higher in samples collected from ccRCC tumor biopsies than it is in non-tumor kidney tissue, whereas *BMAL1* expression in other tumor types is either reduced or unchanged from normal samples of the same tissue type (Figs. 1C and S1). Increased *BMAL1* in ccRCC compared to non-tumor biopsies remains statistically significant when only adjacent samples from the same patients are included in the analysis, suggesting that elevated *BMAL1* in ccRCC samples is not an artefact of tissue collection time or differential sample processing (Fig. S1B). *ARNT2* expression is reduced in ccRCC; *ARNT* and *BMAL2* are unchanged in ccRCC compared to adjacent kidney biopsies from the same patients (Fig S1B).

Correlated expression of twelve genes that are strongly driven by circadian rhythms has been established as a readout of circadian robustness ^1^. Using this measure, we find that circadian rhythmicity is not disrupted in ccRCC, in contrast to other tumor types examined, including papillary RCC, a distinct form of renal cancer that is not driven by HIF2α (Figs. 1D and S2). Data from the Cancer Dependency Map (DepMap) ^29,30^ shows that deletion of *BMAL1* reduces survival of RCC cells (Fig. 1E), indicating that BMAL1 supports growth and/or survival of RCC cells. Genes that act in concert to support cell growth often exhibit correlated dependencies ^31^. In RCC-derived cell lines, dependencies for *ARNT* and *EPAS1* are strongly correlated as expected based on their well-established heterodimeric activation of HIF2α-dependent gene expression. Notably, dependencies for *BMAL1* and *EPAS1* are also strongly correlated in RCC cell lines, while no such correlations are detected for the dependencies of *ARNT2* or *BMAL2* with *EPAS1* dependency across RCC cell lines (Figs. 1F and S3). Together these data suggest that BMAL1 supports the activity of HIF2α in ccRCC.

### BMAL1 forms an active heterodimer with HIF2α

We and others have shown that BMAL1 can interact with HIF1α ^23–25,32^. To evaluate the potential for BMAL1 to partner with HIF2α in ccRCC cells, we expressed FLAG-tagged BMAL1 in 786O cells. 786O cells were established from a ccRCC tumor biopsy in which VHL is inactive, resulting in constitutive stabilization of endogenous wildtype HIF2α. By immunoprecipitation of the FLAG tag, we found that BMAL1 interacts with endogenous HIF2α (Fig. 1G). To evaluate whether BMAL1 can cooperate with HIF2α to activate target gene expression, we used a luciferase reporter under the control of a hypoxia response element derived from the PGK1 promoter region (*HRE-Luciferase*). We demonstrate that overexpression of either ARNT or BMAL1 enhances activation of *HRE-Luciferase* by HIF2α (Fig. 1H). Similar to previous reports describing their transactivation of HIF1α^24,32^, BMAL1 confers greater transcriptional activity than ARNT does.

To determine whether BMAL1 can form a stable complex with HIF2α *in vitro*, we co-expressed and purified the two proteins from insect cells. BMAL1 and HIF2α co-eluted during heparin chromatography, and SDS-PAGE analysis indicated that they formed a stoichiometric complex (Fig. 2A). Analysis by mass photometry of the purified sample (Fig. 2B) further confirmed that the two proteins formed a stable heterodimeric complex, even at low concentration (20 nM).

**Figure 2:**
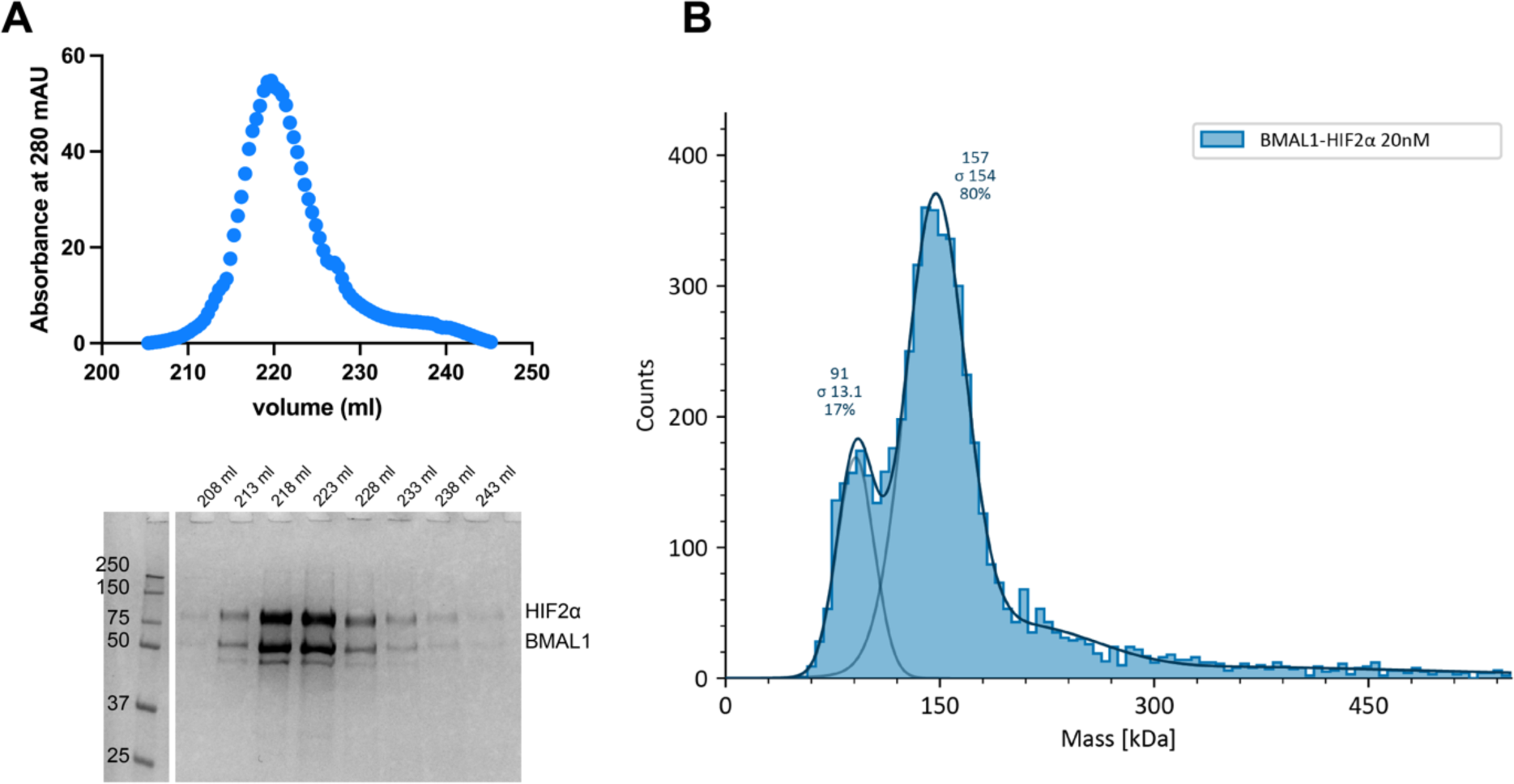
Purified BMAL1 and HIF2α form a stable complex in vitro. (**A**) Heparin chromatography elution of BMAL1 and HIF2α co-expressed in insect cells. SDS-PAGE analysis shows a co-eluted stoichiometric complex of BMAL1-HIF2α. (**B**) Mass photometry of purified BMAL1-HIF2α complex. A minor peak centered at 91 kDa corresponds to the molecular weight of HIF2α, suggesting that it is in slight excess. The major peak, centered at 157 kDa, is consistent with the calculated molecular weight for the BMAL1-HIF2α heterodimer.

### BMAL1 regulates HIF2α target gene expression in ccRCC cells

To measure the contributions of endogenous ARNT and BMAL1 to HIF2α-driven gene expression in ccRCC cells, we sequenced RNA prepared from 786O cells in which either ARNT or BMAL1 was depleted by shRNA. Efficient depletion of BMAL1 or ARNT was confirmed by Western blot (Fig. 3A). We used DESeq2 ^33^ to identify transcripts that were significantly altered and found a striking overlap between the genes affected by loss of ARNT and those affected by loss of BMAL1 (Fig. 3B). Because HIF2α and BMAL1 are expected to primarily activate the expression of their transcriptional targets, we focused on genes that exhibit significantly decreased expression upon depletion of ARNT or BMAL1 as more likely direct targets (Fig. 3C): 42.7% or 54.3% of the transcripts that were significantly decreased by *shBMAL1* or by *shARNT* were decreased by both shRNAs (Fig. 3C,D). Hallmark gene sets generated from multiple primary experiments represent the transcripts regulated by pathways of interest with high confidence across experimental conditions ^34^. We examined the expression of 200 transcripts in the Hallmark HYPOXIA gene set ^34^ using gene set enrichment analysis (GSEA) ^35^ and found that they are robustly impacted by depletion of either *ARNT* or *BMAL1* (3E-G). In a separate experiment, we used 786O cells expressing wildtype VHL (WT8 cells) to highlight transcripts impacted by VHL-dependent suppression of HIF2α. Notably, all transcripts altered by depletion of *BMAL1* in 786O cells were also affected by rescue of VHL (Fig. 3H and S4).

**Figure 3.**
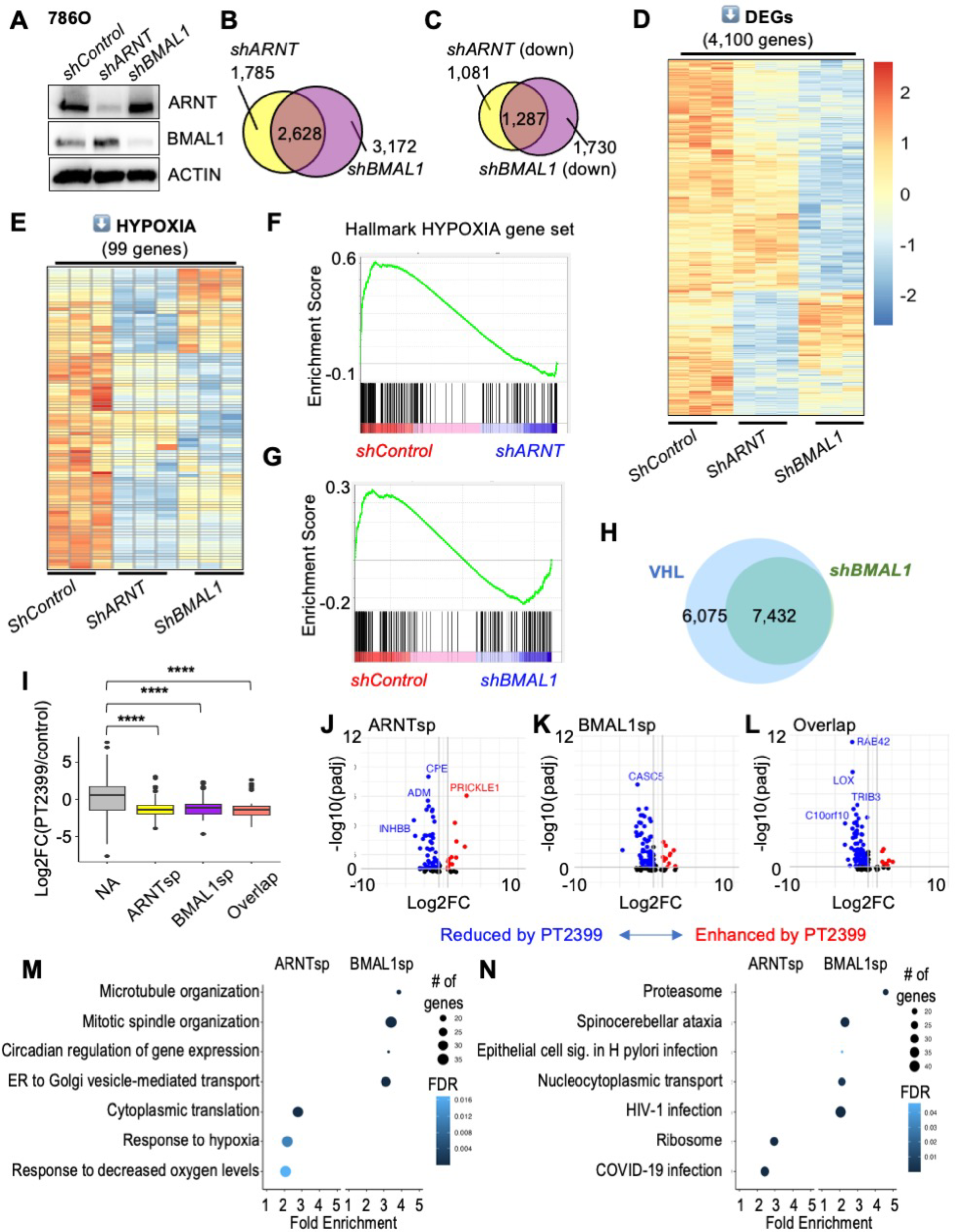
Endogenous BMAL1 contributes to HIF2α target gene expression in RCC cells. **(A)** Detection of ARNT, BMAL1, and ACTIN by immunoblot in 786O cells expressing the indicated shRNAs. (**B-E**) Venn diagrams (*B,C*) and heatmaps (*D,E*) depicting all differentially expressed genes (DEGs) (*B*), significantly downregulated genes (*C,D*) or downregulated genes in the Hallmark HYPOXIA gene set (*E*) in 786O cells expressing the indicated shRNAs. DEGs were identified using DESeq2 with a false discovery rate (FDR) cutoff of 0.1. (**F,G**) Enrichment plots showing the impact of *shARNT* (*F*) or *shBMAL1* (*G*) on genes in the Hallmark HYPOXIA gene set. (**H**) Venn diagram depicting overlap of DEGs in 786O cells expressing VHL (WT8 cells) or expressing *shBMAL1*. (**I**) Boxplot depicting changes in gene expression in PDXs treated with PT2399 (data from ^26^ including sensitive PDXs only) for genes grouped by whether their expression in 786O cells is decreased by *shARNT* and not by *shBMAL1* (ARNTsp, yellow), by *shBMAL1* and not by *shARNT* (BMAL1sp, purple), by either *shARNT* or *shBMAL1* (Overlap, salmon), or neither (NA, gray). **** P < 0.0001 by two-way ANOVA with Tukey’s correction. Boxes depict the median and interquartile range (IQR), whiskers extend either to the minimum or maximum data point or 1.5*IQR beyond the box, whichever is shorter. Outliers (values beyond the whisker) are shown as dots. (**J-L**) Volcano plots depicting expression changes for individual genes in groups depicted in (*I*). Genes with padj < 0.05 are colored in red (fold change > 1.5) or blue (fold change < 0.67). (**M,N**) Top non-redundant GOBP (*M*) or KEGG (*N*) pathways with ≥ 15 genes, FDR < 0.05, fold enrichment ≥ 2 enriched among ARNT-specific or BMAL1-specific target genes in 786O cells.

We took advantage of data from a previous study ^26^ that examined the impact of the HIF2α antagonist drug PT2399 on gene expression in patient-derived xenograft (PDX) tumors to ask how transcripts that are specifically dependent on either ARNT or BMAL1 are affected by disruption of HIF2α heterodimers *in vivo*. We find that genes that are reduced by depletion of ARNT and/or BMAL1 in 786O cells exhibit significantly lower expression in PDX samples that are sensitive to growth inhibition by PT2399 when treated with the drug compared to those treated with vehicle alone (Fig. 3I-L). A more detailed analysis reveals that ARNT-specific targets are enriched in genes related to hypoxia response, ribosome, and metabolism pathways and have higher GC content and BMAL1-specific targets are enriched in genes related to mitosis, intracellular transport, proteasome and circadian rhythm pathways and have greater transcript lengths, more exons, and longer 5’ untranslated regions (Fig. 3M,N and S5). Similar outcomes were observed upon depleting ARNT or BMAL1 in A498 cells (Fig. S6). Together, these findings show that endogenous ARNT and BMAL1 regulate the expression of overlapping and distinct HIF2α target genes in ccRCC patient-derived cells.

### BMAL1 influences HIF2α recruitment to chromatin

HIF2α promotes transcription as part of a heterodimeric complex that interacts with hypoxia response elements (HREs: 5’-N(G/A)CGTG-3’), which are closely related to the canonical E-box sequence bound by BMAL1-CLOCK heterodimers (5’-CACGTG-3’). This suggests that BMAL1 could influence HIF2α target gene expression through diverse mechanisms, including transcriptional activation by BMAL1-HIF2α heterodimers and competition for sites that match both recognition sequences. Subsets of target sites are likely preferentially regulated by alternate bHLH-PAS heterodimers with distinct sequence preferences. To characterize the localizations of endogenous BMAL1 and HIF2α in native chromatin and how these are impacted by depletion of BMAL1, we sequenced genomic DNA associated with BMAL1 or HIF2α in 786O cells. We used MACS2 ^36^ to identify 1,813 and 1,204 genomic regions enriched in chromatin purified with BMAL1 or HIF2α, respectively (Fig. 4A,B). Consistent with prior reports ^37,38^, genomic regions associated with BMAL1 and HIF2α are enriched in promoters and introns (Fig. S7A). 336 loci were identified as co-occupied by BMAL1 and HIF2α, representing 18.5% or 27.9% of the sites associated with BMAL1 or HIF2α, respectively (Fig. 4A). We used Hypergeometric Optimization of Motif EnRichment (HOMER) ^39^ to identify sequence motifs that are enriched in chromatin associated with HIF2α and/or BMAL1. The top motifs identified include those that have been defined biochemically to be preferentially associated with bHLH-PAS transcription factors, including CLOCK, BMAL1, ARNT, HIF1α, and HIF2α (Fig. 4C). Notably, motifs associated with CLOCK and NPAS were enriched uniquely in the BMAL1 cistrome, while the HIF-1b motif was identified only in the HIF2a cistrome (Fig. 4C). These findings provide confidence in the sensitivity and specificity of the data and demonstrate that BMAL1 and HIF2α co-occupy a sizeable fraction of each of their cistromes in 786O ccRCC cells.

**Figure 4.**
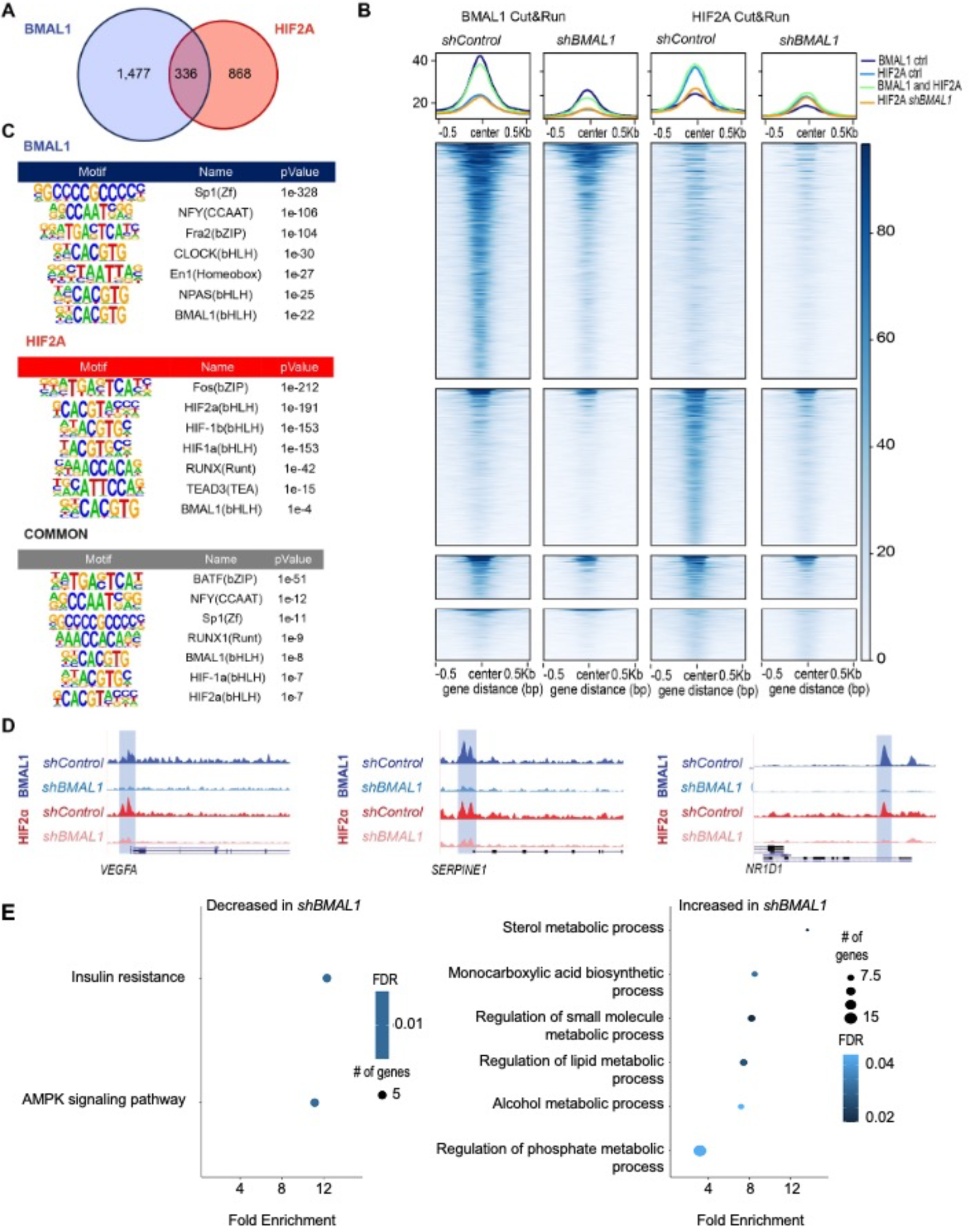
BMAL1 influences recruitment of HIF2α to a subset of target genes. (**A**) Venn diagram depicting the numbers of genomic sites (“peaks”) identified in chromatin fragments isolated by CUT&RUN procedure from 786O cells using antibodies recognizing BMAL1 (blue) or HIF2α (red). **(B**) Chromatin binding profiles of BMAL1 and HIF2α in CUT&RUN samples (n=3 per condition) prepared from 786O cells expressing the indicated shRNAs. Peaks are depicted in four groups: BMAL1 peaks in 786O cells expressing *shControl* (top row: 1,813 peaks), HIF2α peaks in 786O cells expressing *shControl* (second row: 1,207 peaks), peaks associated with both BMAL1 and HIF2α in 786O cells expressing *shControl* (third row: 336 peaks), or HIF2α peaks identified only in 786O cells expressing *shBMAL1* (bottom row: 393 peaks). (**C**) Transcription factor binding motifs enriched in chromatin associated with BMAL1, HIF2α, or both (common) in *shControl* cells. (**D**) Representative genome browser tracks for BMAL1 and HIF2α CUT&RUN in 786O cells expressing *shControl* or *shBMAL1*, showing peaks in *VEGFA*, *SERPINE1*, and *NR1D1* loci. Data represent merged read counts for triplicate samples for each condition. (**E,F**) Combined KEGG and GOBP pathways enriched (≥ 5 genes, FDR < 0.05, fold enrichment > 1.5) in genes located near peaks identified in both BMAL1 and HIF2α CUT&RUN samples (336 common peaks) and exhibiting significantly decreased (*E,* 1,730 genes) or increased (*F,* 1,442 genes) expression in 786O cells expressing *shBMAL1*. This analysis integrates CUT&RUN data with RNA sequencing data described in Figure 3.

Depletion of BMAL1 reduced chromatin association of both BMAL1 and HIF2α at many sites that were occupied in control cells and reduced the number of significantly enriched loci detected in chromatin purified with BMAL1 (Fig. 4B, 4D and S7B). Although MACS2 indicated several peaks bound to HIF2α exclusively in BMAL1-depleted 786O cells, visual inspection and motif enrichment analyses do not support widespread redistribution of HIF2α in BMAL1-depleted cells (Figs. 4B, S7C-E). Instead, HIF2α seems to be absent from a subset of its target loci in BMAL1-depleted 786O cells and its association with other genomic regions is preserved. By integrating CUT&RUN results with RNA sequencing, we found that genes near chromatin loci bound to both BMAL1 and HIF2α that exhibited significantly altered RNA expression in BMAL1-depleted cells are enriched in pathways related to metabolic functions (Fig. 4E). Genes that are associated with HIF2α in control cells and exhibit enhanced expression upon BMAL1 depletion are enriched in pathways related to angiogenesis (Fig. S7E). Together, these data indicate that BMAL1 and HIF2α co-occupy a subset of the canonical target sequences for each of them in native chromatin, and suggest that BMAL1 may preferentially promote expression of HIF2α target genes that impact metabolism over those related to vascular remodeling.

### Depletion of *BMAL1* suppresses growth in RCC cells *in vitro* and *in vivo*

We measured clonogenicity to investigate whether BMAL1 promotes growth of ccRCC cells (786O, RCC4, and A498) and found that depletion of *BMAL1* reduces colony formation in cells plated at low density (Fig. 5A-C). To evaluate the impact of BMAL1 *in vivo*, we generated xenograft tumors in immunocompromised murine hosts. Depletion of *BMAL1* suppressed the growth of tumors derived from 786O or A498 cells *in vivo* (Fig. 5D). Clinically, ccRCC is much more common in men than in women ^5^. Here, we observed no difference in the growth of cell-derived xenograft tumors implanted in male or female hosts (Fig. 5D), suggesting that factors contributing to sexual dimorphism in ccRCC are not present in this model system.

**Figure 5.**
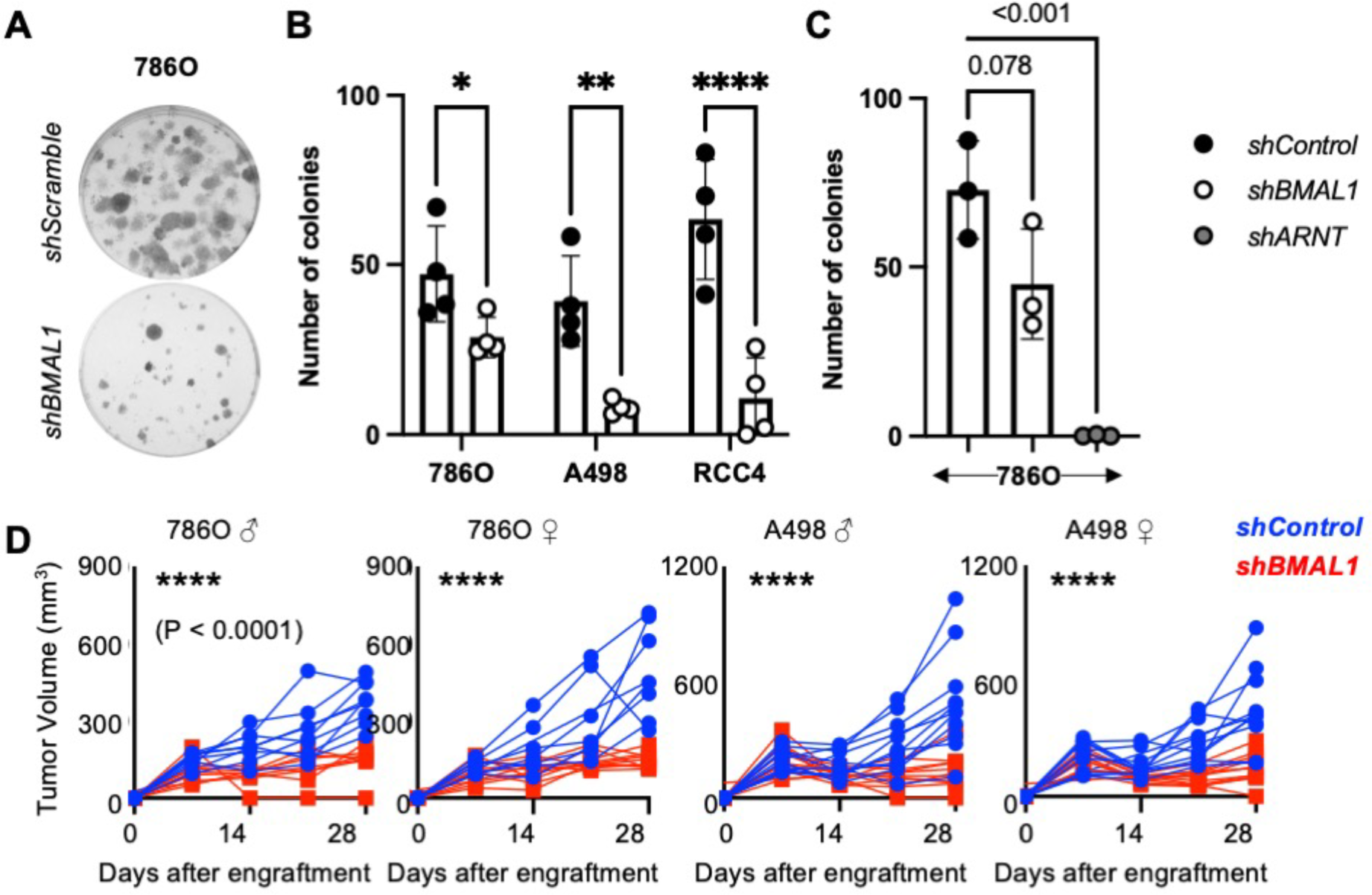
Depletion of *BMAL1* suppresses growth in RCC cells and tumors. (**A-C**) Representative images (*A*) and quantification (*B,C*) of colonies stained with crystal violet 10-16 days after plating 250 cells expressing the indicated plasmids per well. Data represent the mean ± s.d. for 3-4 wells per condition. * P < 0.05, ** P < 0.01, *** P < 0.001 by two-way ANOVA with Tukey’s correction for multiple hypothesis testing. (**D**) Volume of xenograft tumors grown in flanks of male or female NIH-III Nude mice from implanted 786O or A498 cells expressing indicated shRNAs. Weekly measurements of individual tumor volumes are shown. **** P < 0.0001 for *shBMAL1* vs *shControl* by repeated measures ANOVA.

### *BMAL1* enhances growth suppression by HIF2α antagonists

Based on the critical requirement for HIF2α to drive the formation and growth of ccRCC, elegant work led to the development of HIF2α antagonists that disrupt ARNT-HIF2α heterodimers by interacting with a surface pocket in the PAS-B domain of HIF2α ^26^. HIF2α antagonists are effective at reducing the growth of many ccRCC tumors, but resistance to these drugs in up to 30% of cases remains unexplained ^14,26,40^. We analyzed publicly available RNA sequencing data from a study that investigated differences between patient-derived xenograft tumors that were either sensitive or resistant to growth suppression by HIF2α antagonists ^26^. *BMAL1* mRNA expression was higher in patient-derived xenografts that were sensitive to growth suppression by PT2399 (Fig. 6A). HIF2α antagonists like PT2399 disrupt ARNT-HIF2α by causing a conformational change in HIF2α PAS-B resulting in a clash between a methionine (M252) in HIF2α and a glutamine (Q447) in ARNT ^28^. BMAL1 contains a similarly bulky amino acid (M423) in the analogous loop of the PAS-B domain (Fig. S8A), suggesting that BMAL1-HIF2α heterodimers would also be disrupted by HIF2α antagonists. To evaluate whether BMAL1-HIF2α is disrupted by HIF2α antagonists, we expressed a stabilized mutant HIF2α with FLAG-tagged ARNT, ARNT2, BMAL1, or BMAL2 in HEK293 cells. Purification of FLAG-tagged proteins revealed that the interactions of HIF2α with each of these partners are disrupted by PT2399, with BMAL1-HIF2α appearing to be more readily disrupted than ARNT-HIF2α (Fig. 6B). To quantitatively compare the impact of PT2399 on the transactivation activities of ARNT-HIF2α and BMAL1-HIF2α, we turned to luciferase reporter assays. Strikingly, expression of HRE-driven luciferase is much more sensitive to suppression by PT2399 in cells overexpressing stabilized HIF2α in combination with BMAL1 than it is in cells in which stabilized HIF2α is combined with overexpression of ARNT (Fig. 6C).

**Figure 6.**
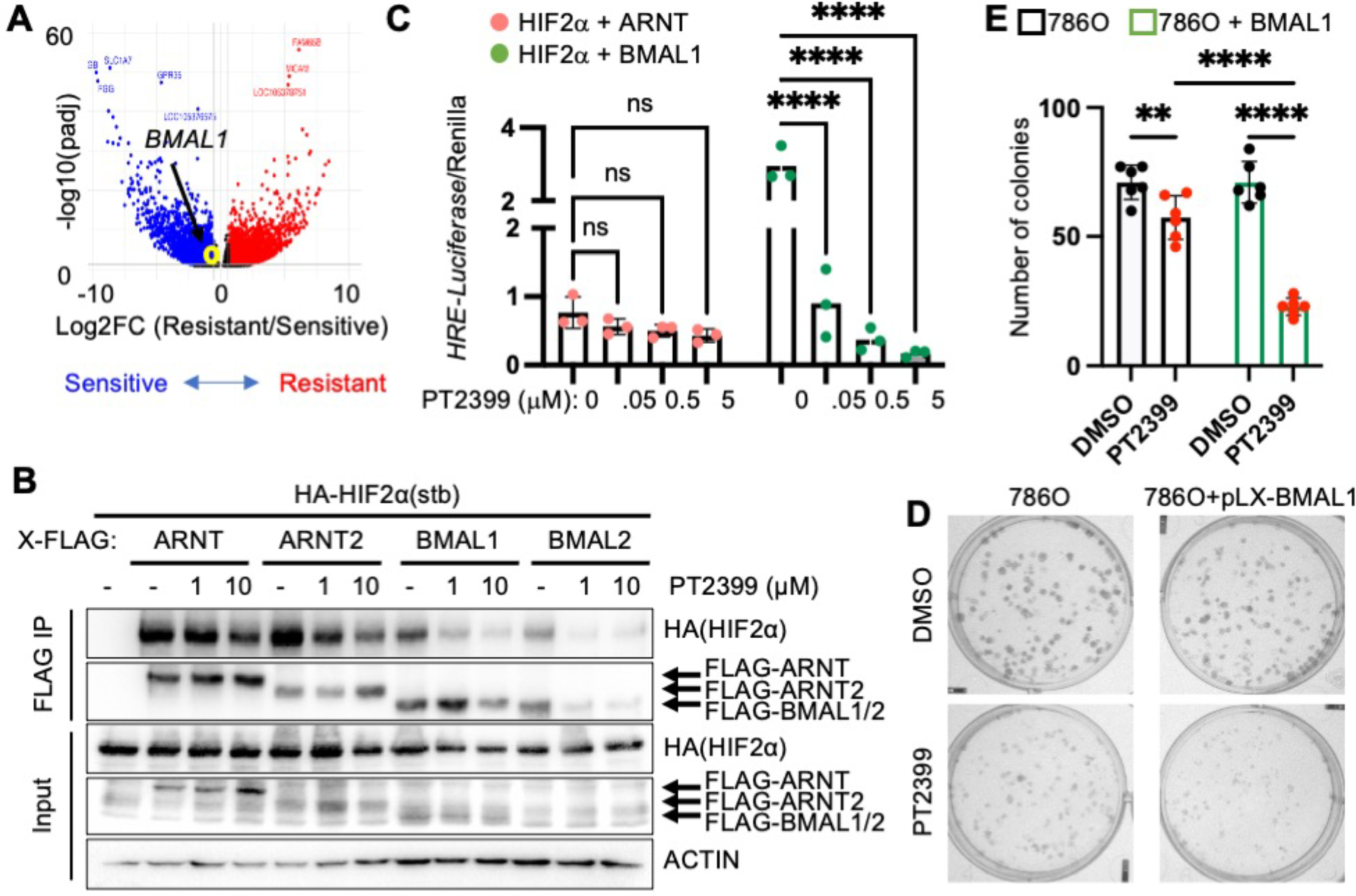
*BMAL1* promotes sensitivity to PT2399. (**A**) Volcano plot depicting differentially expressed genes in patient-derived xenografts that were sensitive or resistant to growth suppression by PT2399 in ^26^. Genes with padj < 0.05 are colored in red (FC > 1.5) or blue (FC < 0.67). (**B**) Detection of indicated proteins in cell extracts (input) or following immunoprecipitation of the FLAG tag from HEK293 cells transiently expressing indicated plasmids HIF2α(stb): stabilized HIF2α (P405A, P531A, N837A) and treated with 10 μM MG132 for 4 hours and indicated concentrations of PT2399 for 1 hour. (**C**) Relative luciferase units detected from U2OS cells expressing *HRE-Luciferase* and additional indicated plasmids and treated with PT2399 at indicated concentrations for 16 hours. P < 0.0001 for interaction between bHLH partner and PT2399 treatment. (**D,E**) Representative images (*D*) and quantification (*E*) of colonies stained with crystal violet 10-16 days after plating 250 cells expressing the indicated plasmids per well in media containing vehicle (DMSO, black circles) or 5 μM PT2399 (red circles). Bars with black and green outlines represent 786O cells with or without overexpression of BMAL1, respectively. In (*C*) bars represent mean ± s.d. for three independent experiments and symbols represent the mean of n=5 measurements for each experiment. Data in (*E*) represent the mean ± s.d. for 3-6 samples per condition from one experiment representative of at least three replicates. ** P < 0.01, **** P < 0.0001 by two-way ANOVA with Sidak’s (*C*) or Tukey’s (*E*) correction for multiple hypothesis testing.

To investigate the functional impact of the observed sensitivity of BMAL1-HIF2α to HIF2α antagonist treatment on cell growth, we used lentivirus to overexpress BMAL1 in 786O cells. Using this approach, we found that increased expression of BMAL1 enhanced the sensitivity of 786O cells to growth inhibition by the HIF2α antagonist PT2399 (Fig. 6D,E and S8B-D). Together, these findings suggest that BMAL1 could play an important role in determining the sensitivity of ccRCC to HIF2α antagonist drugs.

## Discussion

The circadian transcription factor BMAL1 is closely related to ARNT, the canonical partner for HIF alpha subunits. We demonstrate here that BMAL1 directly participates in HIF2α target gene regulation and promotes growth in ccRCC-derived cells and xenograft tumors. Interaction between BMAL1 and a highland-adapted variant of HIF2α influences circadian rhythms in a Tibetan rodent ^41^, and BMAL1-HIF2α heterodimers can contribute to circadian rhythms in myocardial injury ^42^. These findings provide additional support for the idea that BMAL1 is an important partner in HIF2α-driven gene regulation. Circadian disruption enhances the risk of several cancer types ^43^ and deletion of *BMAL1* has been used to study circadian disruption genetically, with mixed results in mouse models of cancer ^2–4,44^. The reasons for diverse impacts of *BMAL1* deletion in different tumor types are unclear. Our findings suggest that decreased expression of HIF target genes could contribute to reduced growth upon *BMAL1* deletion in some tumor types.

Two additional homologs, ARNT2 and BMAL2, could participate in HIF2α signaling in a similar manner. We found that *ARNT2* expression is significantly reduced and *BMAL2* is enhanced in ccRCC samples compared to adjacent normal kidney tissue (Fig. S1B). A pre-publication report indicates that BMAL2 supports hypoxic responses in a pancreatic cancer model ^45^, so it could also play a role in ccRCC. Further investigation is needed to understand the contributions of diverse bHLH-PAS partners to HIF2α activities and responses to HIF2α antagonist drugs in diverse physiological and pathological contexts.

HIF1α and HIF2α activate gene expression through HREs and regulate overlapping and distinct sets of target genes. Differences in their transcriptional targets are presumed to underlie the greater dependence of ccRCC on HIF2α activity, but the determinants of differential specificity are unclear. Depletion of either *ARNT* or *BMAL1* in ccRCC-derived cell lines dramatically altered gene expression, including that of hypoxia target genes. Some HIF targets were reduced upon depletion of *BMAL1* and others were enhanced, suggesting that subsets of genes are preferentially activated by distinct HIF2α-containing heterodimers. This possibility is further supported by enrichment of overlapping and distinct nucleotide sequences in chromatin purified with BMAL1 or HIF2α from ccRCC-derived cells. Despite the divergent impacts on gene expression of depleting ARNT or BMAL1 in ccRCC patient-derived cell lines, losing either of these HIF2α partners dramatically reduces the ability of several ccRCC cell lines to form colonies *in vitro* and xenograft tumors *in vivo*. Additional investigation will be required to determine whether specific HIF2α target genes critical for tumor growth require both ARNT and BMAL1 to reach an expression threshold that is needed to support tumor formation or if loss of distinct genes driven by ARNT-HIF2α or by BMAL1-HIF2α contribute to growth impairment upon depletion of each heterodimer. Notably, direct interaction between two BMAL1-CLOCK heterodimers and histones can promote gene expression via tandem E-boxes and the BMAL1-CLOCK heterodimer was shown to compete with histones for DNA access ^22^. These observations suggest mechanisms by which multiple bHLH-PAS heterodimers could cooperatively influence the expression of common target genes. Additional research is needed to determine how ARNT and BMAL1 cooperate to support HIF2α activities in ccRCC.

Approximately 30% of ccRCC patient-derived xenograft tumors are resistant to HIF2α antagonists, with resistant tumors exhibiting no significant changes in gene expression following treatment ^26^. Thus, sensitivity of ccRCC to treatment with HIF2α antagonists is associated with changes in gene expression; and such sensitivity has been shown to require HIF2α ^28^. There is currently a lack of comprehensive understanding regarding mechanisms underlying resistance to HIF2α antagonists^14,26–28^. Here, we showed that *BMAL1* expression is higher in PDXs that were sensitive to growth inhibition by PT2399 and that HIF2α-BMAL1 heterodimers are more sensitive to suppression by PT2399 than HIF2α-ARNT heterodimers are. We defined groups of genes as ARNT-specific or BMAL1-specific by RNA sequencing of ccRCC patient-derived cell lines in which ARNT or BMAL1 is depleted by shRNA. Expression for both groups was reduced by PT2399 treatment in patient-derived xenografts in which PT2399 is effective at suppressing *in vivo* tumor growth, further supporting the idea that ARNT and BMAL1 promote the expression of overlapping and distinct sets of HIF2α target genes that are relevant for therapeutic responses to HIF2α antagonists in ccRCC. Finally, increasing BMAL1 expression in ccRCC-derived cells rendered them more sensitive to growth inhibition by PT2399. Together, these findings suggest that BMAL1 enhances sensitivity to HIF2α antagonists through the formation of a BMAL1-HIF2α heterodimer that is more sensitive to suppression by HIF2α PAS-B domain ligands.

## Methods

### Analyses of RNA sequencing data from TCGA projects

RNA sequencing data for five projects in The Cancer Genome Atlas (TCGA) and from the clinical proteomic tumor analysis consortium 3 (CPTAC3) were downloaded from the NIH genome data commons (https://portal.gdc.cancer.gov/). Expression of *ARNT*, *ARNT2*, *BMAL1*, and *BMAL2* was extracted, analyzed, and visualized in Rstudio using packages rstatix and ggpubr. Clock correlation distance analysis was performed using the online tool available through the Hughey lab (https://hugheylab.shinyapps.io/deltaccd/). Software used for statistical analysis and data visualization will be available via GitHub.

### Cancer Dependency Map Analysis

Dependency data (DepMap_Public_23Q4+Score,_Chronos) for 37 kidney cell lines were downloaded from the Cancer Dependency Map portal (https://depmap.org/portal/) on February 29, 2024. Statistical analysis and data visualization were performed in Rstudio using packages rstatix and ggpubr. Software will be available at GitHub.

### Cell culture

786-O (ATCC® CRL-1932™), A-498 (ATCC® HTB-44™), HEK293T (ATCC® CRL3216™), and U2OS (ATCC® HTB-96™) cells were purchased from the American Type Culture Collection. 786-O (CRISPR-control), WT8, and RCC4 cells were provided by Dr. Celeste Simon. All cell lines were cultured in Dulbecco’s modified Eagle’s medium + 10% fetal bovine serum (Thermo Fisher Scientific) and 1% penicillin-streptomycin (Gibco), and maintained in an atmosphere containing 5% CO2 at 37°C.

### Generation of cell lines expressing shRNA

To generate cell lines expressing shRNA, lentiviral shRNA constructs encoded in PLKO.1 vectors (Sigma-Aldrich, SHC002 (shControl), TRCN0000003816 (shARNT), shBMAL1 (a gift from Dr. Satchidananda Panda), TRCN0000019097 (independent shBMAL1)) were produced by transient transfection in HEK293T cells. Target cells were infected with lentivirus for 4-6 hours before selection in Dulbecco’s modified Eagle’s medium + 10% fetal bovine serum (Thermo Fisher Scientific) and 1% penicillin-streptomycin (Gibco) containing 2.5 µg/mL puromycin for 1 week. After initial selection, cells were maintained in DMEM containing 1.25 µg/mL.

### Co-immunoprecipitation

Transfections in HEK293T cells were performed using polyethylenimine (PEI; Polysciences Inc #23966-2) following standard protocols. pcDNA3.1-HIF2α-HA(Stb) was a gift from Dr. Carrie Partch. ARNT-FLAG, ARNT2-FLAG, and BMAL1-FLAG in the pTwist CMV Hygro vector were purchased from Twist Bioscience. Cells were treated with MG132 (10 μM) for 1 hour before the addition of vehicle control (0.01% Dimethyl sulfoxide (DMSO)) or the indicated concentration of PT2399 (Thermo Fisher Scientific #501932330) for 1 hour before immunoprecipitation.

786O cells were transfected with the pTwist plasmids previously described using Bioscience Lipofectamine® 2000 DNA Transfection Reagent Protocol.

Cells were lysed using RIPA buffer supplemented with protease (Thermo Scientific #A32953) and phosphatase (Sigma #P5266 and #P0044) inhibitors. Protein levels were quantified using the Pierce BCA Protein Assay Kit (Thermofisher #PI23225) and equilibrated before FLAG tagged proteins were immunoprecipitated using anti-Flag M2 agarose beads (Sigma #A2220).

### Western Blotting

Cell lysates were separated using 8% SDS–polyacrylamide gel (National Diagnostics #EC8901LTR) by electrophoresis (Bio-Rad #1658001) and transferred using the Trans-blot Turbo transfer system (Bio-Rad #17001915). Proteins were detected by standard Western blotting procedures.

Primary antibodies used for Western blotting were anti-HA polyclonal (Sigma #H6908), anti-Flag polyclonal (Sigma #F7425), anti-βActin (Sigma #A1978), anti-HIF2a polyclonal (Novus Biologicals #NB100-122), anti-BMAL1 polyclonal (Abcam #ab93806) and anti-Cry1-CT and anti-Cry2-CT as described (Lamia *et. Al*., 2011), and anti-BMAL1 monoclonal (VWR #102231-824). Secondary antibodies used were Goat Anti-Mouse IgG (H + L)-HRP Conjugate (Bio-Rad #1706516), Goat Anti-Rabbit IgG (H + L)-HRP Conjugate (Bio-Rad #1706515), Goat Anti-Guinea Pig IgG-HRP Conjugate (Sigma #A7289). SuperSignal West Pico PLUS Chemiluminescent Substrate (Fisher scientific #PI34095) or Immobilon Forte Western HRP substrate (Sigma # WBLUF0500). Imaging and quantification were performed using the ChemiDoc XRS+ System (Bio-Rad #1708265) and Image Lab software version 6.1.0 build 7. Proteins detected by immunoblotting were normalized to the housekeeping protein β-ACTIN.

### Luciferase assay

U2OS cells infected with lentivirus expressing shRNA targeting “scramble” control sequence (AddGene Plasmid #1864, deposited by Dr. David Sabatini) or shBMAL1 (a gift from Dr. Satchidananda Panda) were seeded at a density of 15,000 cells per 96-well. Cells were transfected using standard polyethylenimine (PEI) protocols in suspension at time of seeding with 30 ng reporter HRELuc (Addgene #26731, deposited by Dr. Navdeep Chandel); 5 ng BMAL1; 15 ng HIF2a; 5 ng for ARNT; 5 ng Renilla Luciferase (a gift from Dr. Ian MacRae). All plasmid dilutions were prepared fresh immediately before transfection. A media change was performed on the day following transfection, at which time vehicle (DMSO) or PT2399 was added where indicated. The following day luciferase activity was measured using the Dual-Glo® Luciferase Assay System (Promega #E2920) and Infinite® 200 PRO microplate reader (TECAN #30190085).

### Protein expression and purification

Full-length human HIF2α and BMAL1, each with an N-terminal Strep tag, were each cloned into pAC8 vectors for insect cell expression. Recombinant baculoviruses were prepared in the *Spodoptera frugiperda* (sf9) cells using the Bac-to-Bac system (Life Technologies). HIF2α and BMAL1 were co-expressed in *Trichoplusia ni* Hive Five cells by infection of 25 ml each of baculoviruses per 1 L of High Five culture. Cells were harvested 48 hours post-infection and lysed by sonication in a buffer containing 25 mM Tris-HCl pH 8.0, 400 mM NaCl, 5% glycerol, 0.5 mM TCEP, 1 mM MgCl2, 1x protease inhibitor cocktail (Roche Applied Science) and 0.1% Triton X-100. Lysate was clarified by ultracentrifugation at 40K RPM for 30 min. The Supernatant was than loaded onto a gravity column for affinity chromatography containing a Strep-Tactin Sepharose bead slurry (IBA life sciences). The column was washed with a high salt (1M NaCl) buffer followed by low salt (200 mM NaCl) buffer, and then eluted at 200 mM NaCl using 5 mM desthiobiotin. The eluted fractions were then diluted to 100 mM NaCl prior to application on a heparin column (GE Healthcare) and then eluted using a linear salt gradient. Finally, samples were dialyzed to no more than 150 mM NaCl and flash frozen in 5% glycerol and stored at -80°C.

### Mass photometry

Prior to mass photometry measurements, protein dilutions were made in MP buffer (20 mM Tris-HCl pH 8.0, 100 mM KCl, and 0.5 mM TCEP). Data were acquired on a Refeyn OneMP mass photometer. 18 μl of buffer was first added into the flow chamber followed by a focus calibration. 2 μl of protein solution was then added to the chamber and mixed, and movies of 60 seconds were acquired. Each sample was measured at least two times independently (n = 2) and Refeyn Discover 2.3 was used to process movies and analyze molecular masses, based on a standard curve created with BSA and thyroglobulin.

### RNA sequencing and analysis

RNA from 786O, WT8, and A498 cells infected with lentivirus expressing shRNA targeting the indicated transcripts was isolated using RNeasy Mini Kit (QIAGEN #74104) and QIAshredder (QIAGEN #79654). RNA purity was assessed by Agilent 2100 Bioanalyzer and quantified by Thermo Fisher Qubit. Total RNA samples were sent to BGI Group, Beijing, China, for library preparation and sequencing. Reads (paired-end 100 base pairs at a sequencing depth of 20 million reads per sample) were generated by BGISEQ-500. In addition, FASTQ files containing RNA sequencing data from Chen et Al. ^26^ were retrieved from the sequence read archive. FASTQ sequencing files were aligned to the GRCh37 Homo sapiens reference genome using SeqMan NGen 17 software (https://www.dnastar.com/manuals/installation-guide). Assembly results were analyzed and counts data were exported using ArrayStar 17 (https://www.dnastar.com/manuals/installation-guide). Differential gene expression analysis (DESeq2) and gene set enrichment analysis (GSEA) were performed using the online tool Gene Pattern (https://www.genepattern.org) to generated normalized count data and identify differentially expressed genes. The RNA-seq FASTQ files were deposited to GEO. GO term analysis was performed using the online tool ShinyGO (http://bioinformatics.sdstate.edu/go/). Data visualization including Venn diagrams, heat maps volcano plots, and GO term representative plots were generated in RStudio using the packages pheatmap, venneuler, and ggplot2. Software used for data visualization will be available via GitHub named by the figure number and panel designation.

### Cleavage Under Targets & Release Using Nuclease (CUT&RUN)

Chromatin immunoprecipitation followed by high-throughput sequencing (ChIP-seq) were performed using CUT&RUN assay kit (CST #86652) following the manufacturer protocol. 100,000 786O cells infected with lentivirus expressing shRNA targeting “scramble” or *BMAL1* were used for each reaction. The primary antibodies used for immunoprecipitation were 5 ug of rabbit mAb IgG isotype as a negative control (CST #66362), 2 ug of rabbit mAb tri-methyl-lys-4 (CST # C42D8), 1 ug of HIF-2α rabbit mAb (CST #59973), or 2 ug of BMAL1 rabbit mAb (CST #14020). 50 pg of spike-in control DNA (provided in kit) was added to each sample for normalization. DNA purification was performed by phenol/chloroform extraction followed by ethanol precipitation. Next generation sequencing libraries were prepared using DNA Library Prep Kit for Illumina (CST #56795) and Multiplex Oligos for Illumina (CST #29580). Libraries were sent to BGI Group, Beijing, China, for sequencing. Reads (paired end 100 base pairs at a sequencing depth of 20 million reads per sample) were generated by DNBSEQ. The CUT&RUN-seq FASTQ files were deposited to X.

### CUT&RUN data analysis

The bioinformatic analysis was conducted at the HPC cluster located at Helmholtz Zentrum München. Initial processing of raw data involved quality control using Fastqc 0.12.1 from the trim galore suite 0.6.10. Subsequently, reads underwent alignment to both the human genome hg19 and the yeast genome sacCer3 using Bowtie2 2.5.3, with the following parameters: --local - -very-sensitive --fr --dovetail --no-mixed -I 10 -X 700. Alignment files (SAM) were then converted to BAM format, and subjected to filtering, and duplicate reads were removed using samtools 1.6 and sambamba 1.0. Peak calling was performed using MACS2 2.2.9.1, specifying parameters --keep-dup all --max-gap 400 –p 1e-5. Post-peak calling, filtering against the hg19 blacklist was executed using bedtools 2.31.1 with the intersect option. Finally, annotation and motif analysis of the peaks was carried out using HOMER 4.11, using annotatePeaks.pl and findMotifsGenome.pl options with the human genome hg19 reference. Peak functional annotation was directly done by Homer using -go option, or with WEB-based GEne SeT AnaLysis Toolkit -WebGestalt (https://www.webgestalt.org/) to identify gene ontologies and KEGG-related pathways after crossing peaks annotation with RNA-seq data.

Spike-in normalization with the aligned reads was achieved against the yeast genome sacCer3 with deeptools 3.5.5 using bamCoverage –scaleFactor --smoothLength 60 --extendReads 150 – centerReads to produce BigWig files. Spike-in scale factor values were calculated as described in the manufacturer protocol (CST #86652). Profiles and heatmap were obtained by using computeMatrix –referencePoint center after spike-in normalization. BigWig files were uploaded to the UCSC genome browser (https://genome-euro.ucsc.edu/index.html) and tracks were visualized against the human genome hg19.

### Colony formation assay

Cells were plated into six-well plates at 250 cells per well, and medium was changed every two or three days. Cells were washed in PBS, fixed for 10 min with 100% methanol, and stained with 0.05% crystal violet for 20 min. Plates were rinsed in deionized H2O, imaged using the ChemiDoc XRS+ System (Bio-Rad), and quantified using FIJI ImageJ (DOI 10.1186/s12859-017-1934-z).

### Cell line derived xenografts

All murine husbandry and experiments were in regulation with the Institutional Animal Care and Use Committee at the Scripps Research Institute (La Jolla, California) under protocol #10-0019. NIH-III nude mice (Charles River Laboratories) were implanted in each flank with 5 X 10^6^ 786O or A498 cells infected with lentivirus expressing shRNA targeting “scramble” or *BMAL1* were suspended in 1:1 ratio of PBS and Matrigel (Corning #CB-40234). The final volume for injection was 100 uL. Mice were ∼8 weeks old and an equal mix of male and female mice. There were 15-20 mice per experimental group. Tumors were measured weekly by caliper and tumor volume was calculated using the formula V = (π/6)(*Length*)(*Width*^2^). Experimental termination was determined empirically when the first mouse had a tumor measuring 600 mm^3^ at which point mice were euthanized by CO2 inhalation.

## Data Availability

RNA sequencing and CUT&RUN sequencing data were deposited into the Gene Expression Omnibus (GEO) database. *We expect to have an accession number available by June 12, 2024, and the data will be made available prior to publication*.

## Acknowledgements

Data used for analyses described in this manuscript were obtained from: the GTEx Portal on 10/30/2023, the sequence read archive (accession number SRP073253), and data generated by the TCGA Research Network (https://www.cancer.gov/tcga). We thank Marie Pariollaud, Megan Vaughan, Fania Feby Ramadhani, Fabiana Quagliarini, Ben Cravatt, Reuben Shaw, Michael Bollong, and Luke Wiseman for helpful discussions, Lara Ibrahim for assistance retrieving published RNA sequencing data and aligning to the human genome, and Judy Valecko for administrative assistance. K.A.L. is supported by National Institutes of Health grants R01CA211187 and R01CA271500. DAV received funding from the DAAD (German Academic Exchange Service) in the context of the Helmholtz Research School for Diabetes, and NHU received funding from the DFG (German Research Foundation), TRR333 BATenergy (450149205).

## Author Contributions

Conceptualization: RM, KAL

Methodology: RM, MCS, KAL, CS, NT, NHU

Investigation: RM, DGC, KAL, CS, DA

Visualization: RM, KAL, CS, DA

Resources: MCS

Funding acquisition: KAL, NT, NHU

Project administration: KAL, NT, NHU

Supervision: KAL, NT, NHU, CJ

Writing – original draft: KAL, CS

Writing – review & editing: RM, MCS, KAL, CS, NT, DA, CJ, NHU

## Competing interests

The authors declare no competing interests.

Supplementary information is available for this paper.

Correspondence and requests for materials should be addressed to Katja A. Lamia;

## Extended data figure legends

**Figure S1.**
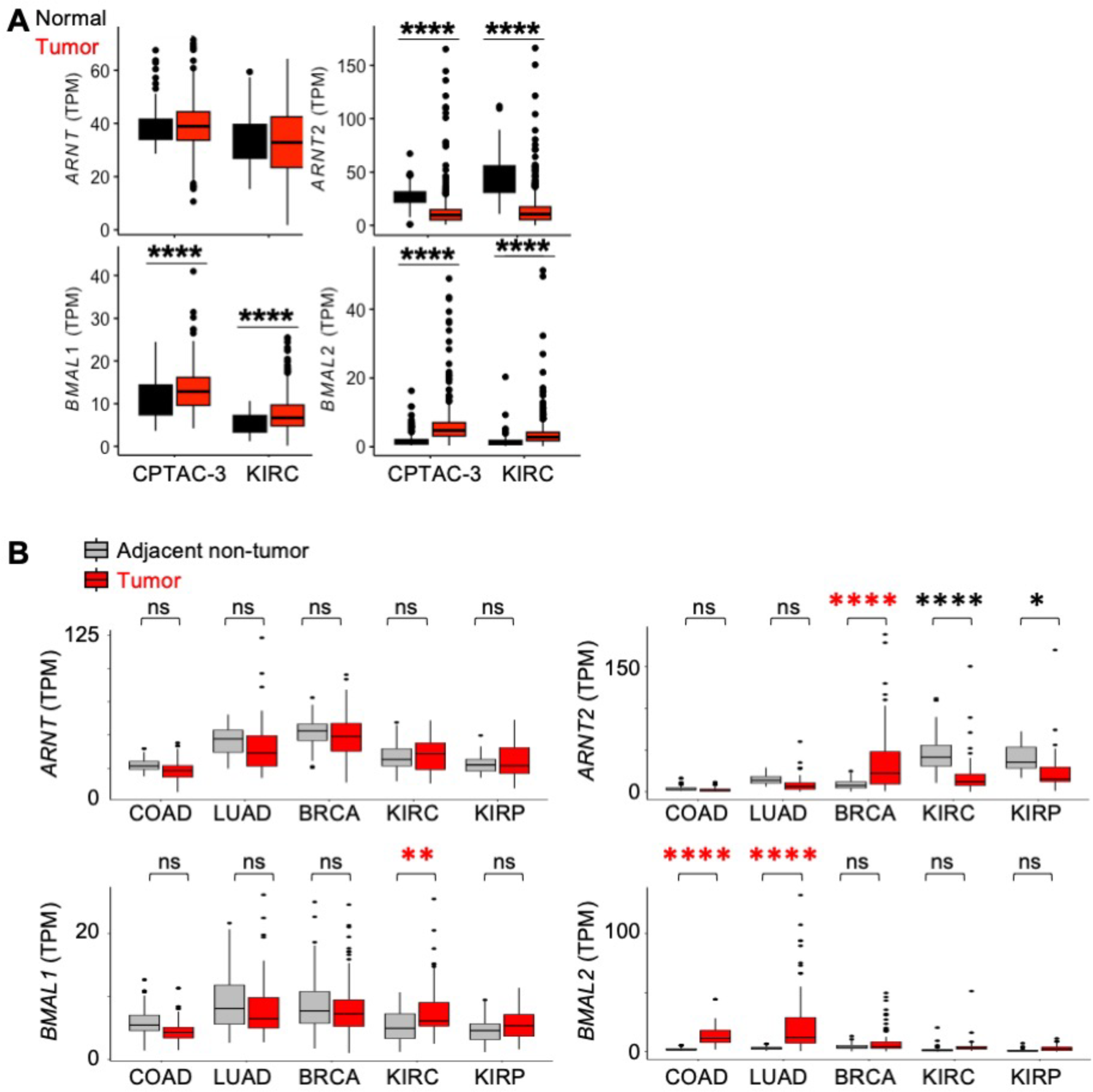
BMAL1 mRNA is elevated in ccRCC. (A,B) Detection of *ARNT*, *ARNT2*, *BMAL1,* and *BMAL2* (transcripts per million, TPM) calculated from RNA sequencing data from ccRCC tumors and normal kidney samples (*A*) or tumors and adjacent normal tissues from the same patients (*B*) in clinical proteomic tumor analysis consortium (CPTAC-3) and cancer genome atlas projects: colorectal adenocarcinoma (COAD), lung adenocarcinoma (LUAD), breast cancer (BRCA), kidney clear cell renal cell carcinoma (KIRC), and renal papillary cancinoma (KIRP). * Padj < 0.05, ** Padj < 0.01, **** Padj < 0.0001 by two-way ANOVA with Tukey’s posthoc correction. In all boxplots, the box depicts the median and interquartile range (IQR), whiskers extend either to the minimum or maximum data point or 1.5*IQR beyond the box, whichever is shorter. Outliers (values beyond the whisker) are shown as dots.

**Figure S2.**
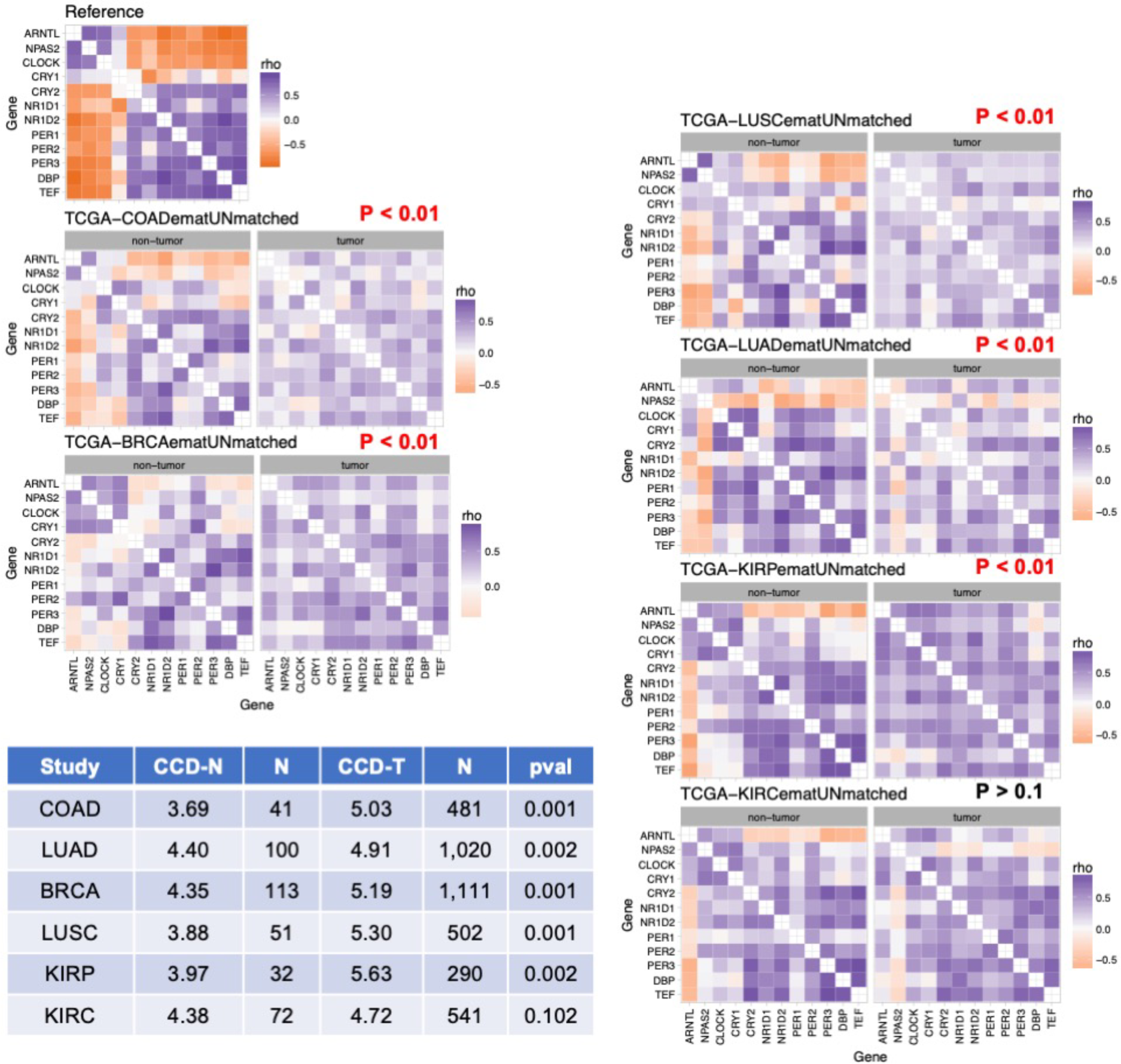
Circadian rhythms are maintained in ccRCC in contrast to other tumor types. Clock correlation distance (CCD) heatmaps calculated from RNA sequencing data from normal human tissues (GTEX, reference) or from tumors and non-tumor samples from the same tissues in cancer genome atlas (TCGA) projects: colorectal adenocarcinoma (COAD), lung adenocarcinoma (LUAD), small cell lung cancer (LUSC), breast cancer (BRCA), kidney clear cell renal cell carcinoma (KIRC), and renal papillary cancinoma (KIRP). The table shows the number of samples in each group (N), calculated CCD values for non-tumor (CCD-N) or tumor (CCD-T) samples, and adjusted p values (pval).

**Figure S3.**
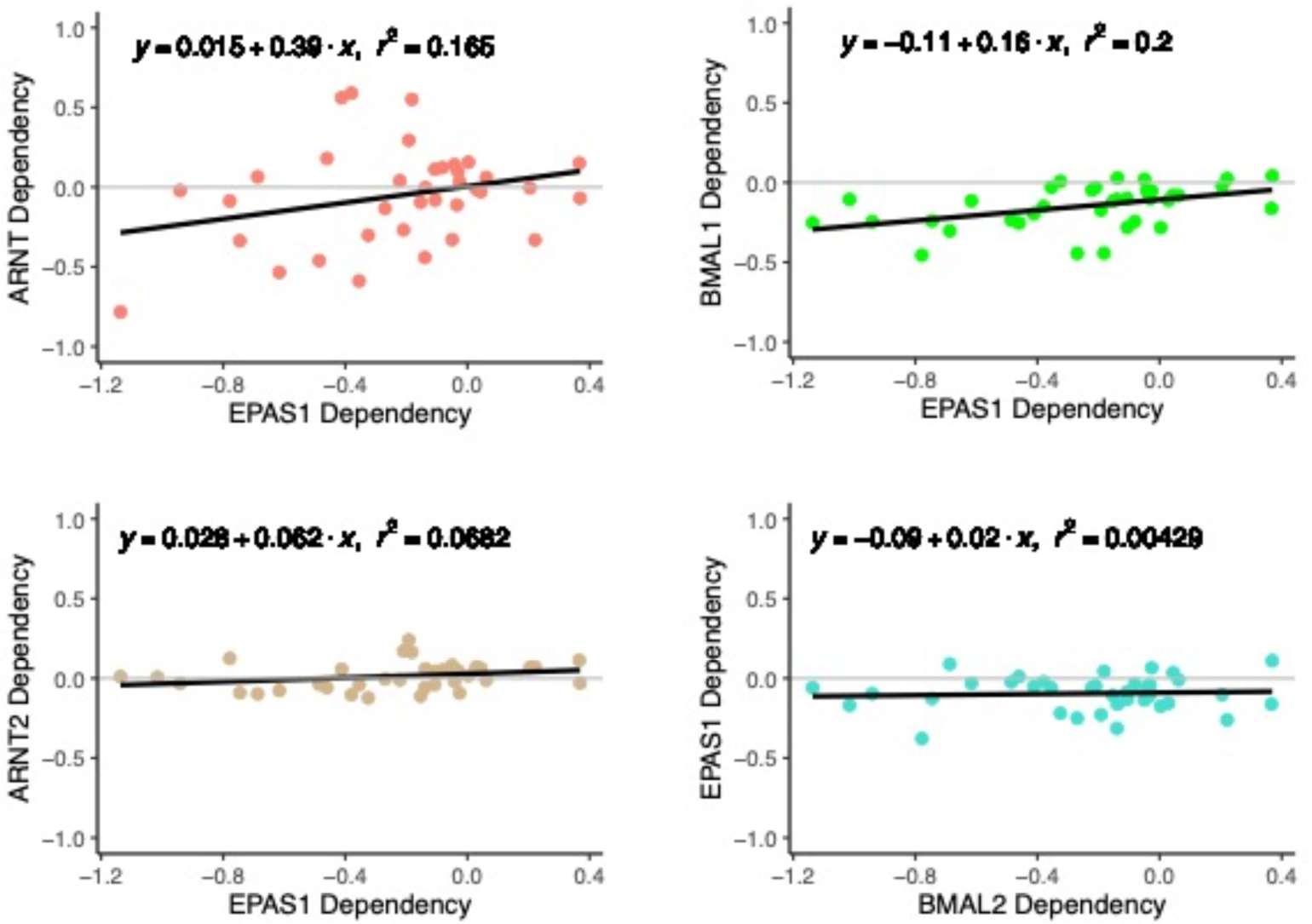
In RCC cell lines, HIF2A (a.k.a. EPAS1) dependency is correlated with dependencies for *ARNT* and *BMAL1*. Correlation of dependency (CHRONOS) scores for *ARNT*, *ARNT2*, *BMAL1*, and *BMAL2* with those for *EPAS1* in RCC cell lines from DepMap ^29,30^.

**Figure S4:**
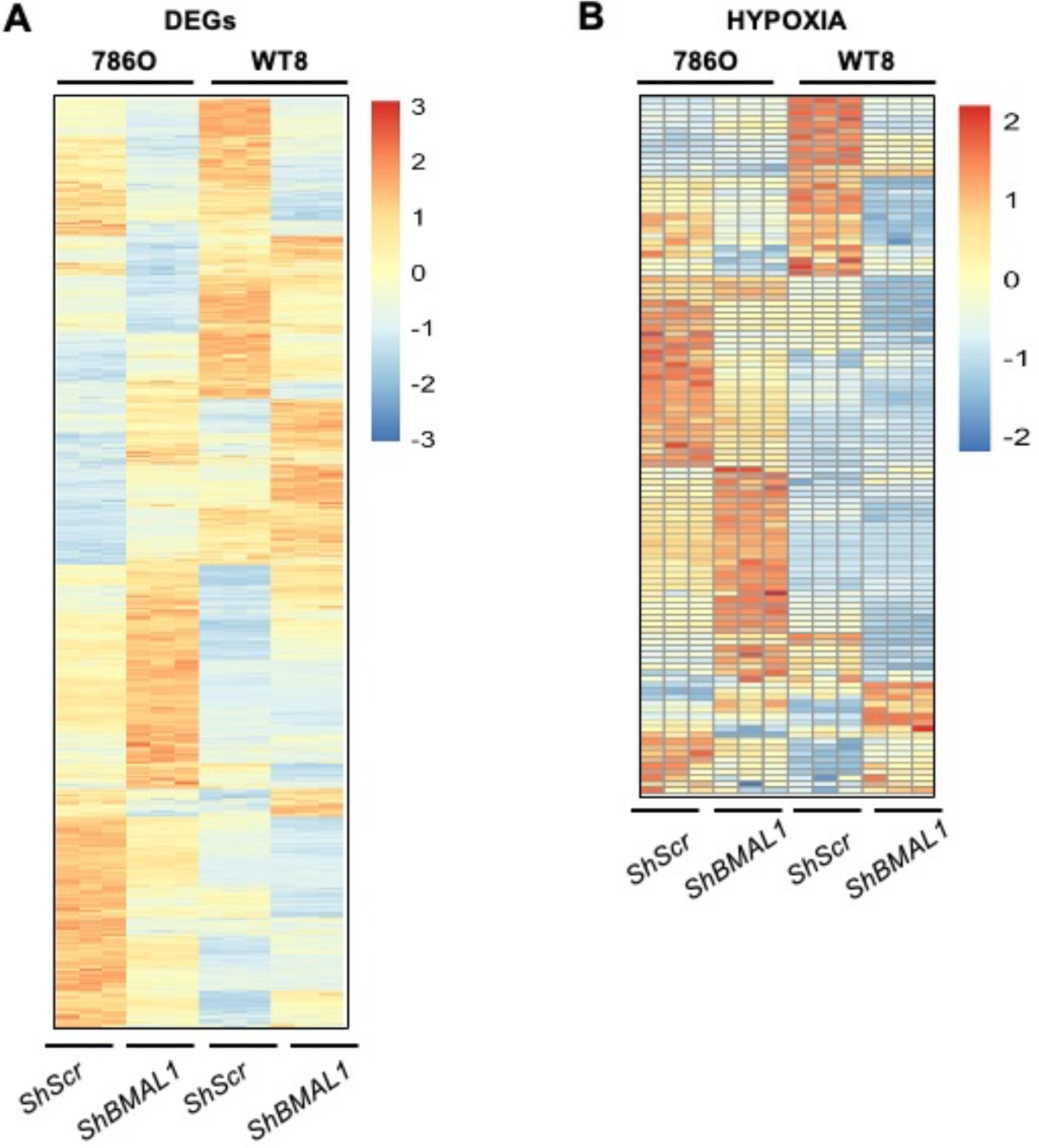
Depletion of BMAL1 alters many of the same genes that are affected by rescuing VHL function in 786O cells. **(A,B)** Heatmaps depicting all differentially expressed genes (DEGs) (*A*), or DEGs in the Hallmark HYPOXIA gene set (*B*) in786O cells and WT8 cells expressing the indicated shRNAs. DEGs were identified using DESeq2 with a false discovery rate (FDR) cutoff of 0.1. (Note: WT8 cells are 786O cells engineered to express WT VHL).

**Figure S5.**
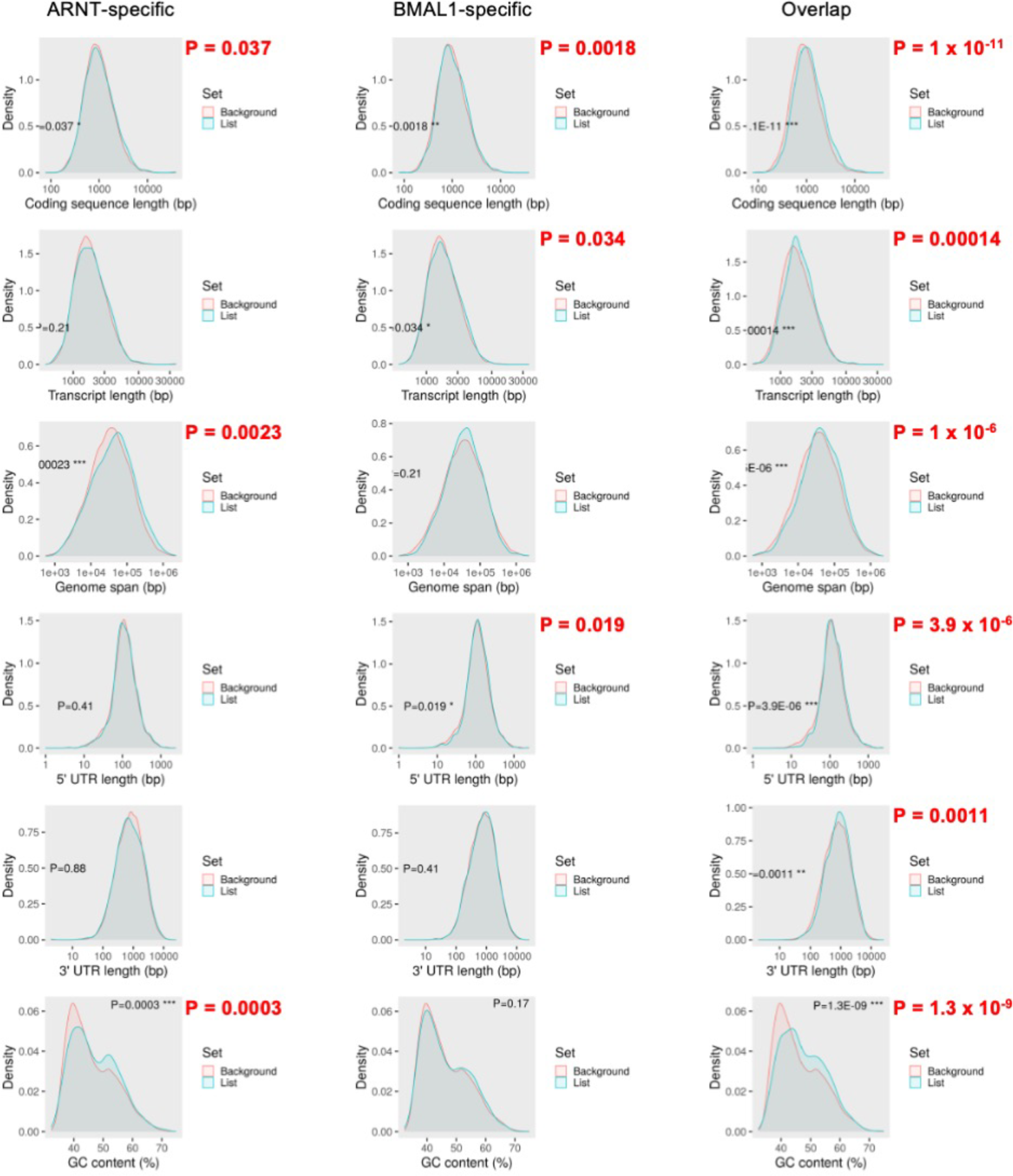
Characterization of ARNT- and BMAL1-dependent transcripts in 786O cells. Distribution of coding sequence length, transcript length, genome span, 5’UTR length, 3’UTR length, and GC content for transcripts that exhibit reduced expression exclusively in 786O cells expressing *shARNT*, exclusively in 786O cells expressing *shBMAL1*, or in both compared to 786O cells expressing *shControl*. Each “List” of DEGs so defined (depicted in blue) is compared to the “background” of all transcripts detected in RNA prepared from 786O cells expressing *shControl* (depicted in red). Analysis performed using ShinyGO ^46^.

**Figure S6.**
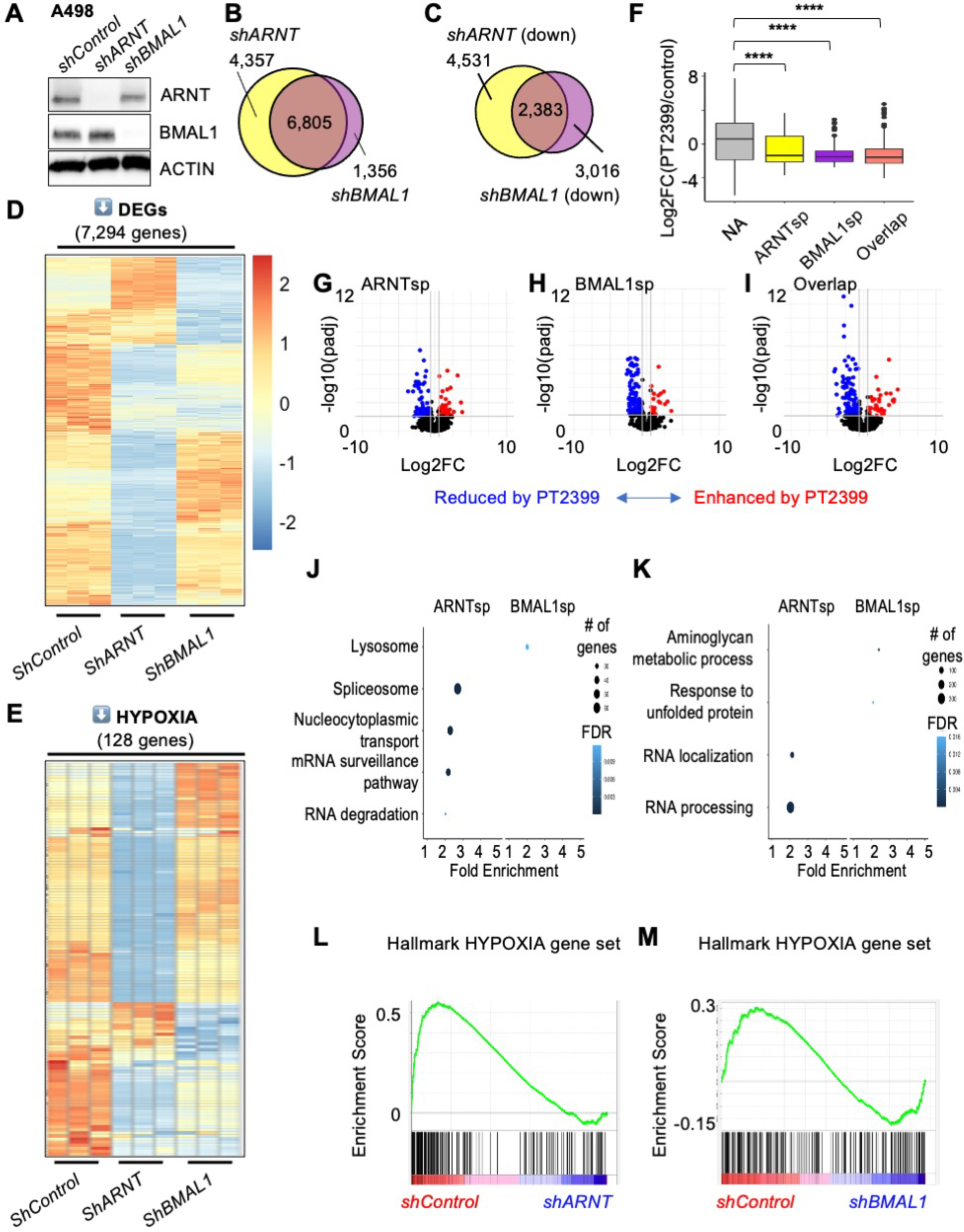
Endogenous BMAL1 contributes to HIF2α target gene expression in A498 cells. **(A)** Detection of ARNT, BMAL1, and ACTIN by immunoblot in A498 cells expressing the indicated shRNAs. (**B-E**) Venn diagrams (*B,C*) and heatmaps (*D,E*) depicting all differentially expressed genes (DEGs) (*B*), significantly downregulated genes (*C,D*) or downregulated genes in the Hallmark HYPOXIA gene set (*E*) in A498 cells expressing the indicated shRNAs. DEGs were identified using DESeq2 with a false discovery rate (FDR) cutoff of 0.1. (**F**) Boxplot depicting changes in gene expression in patient-derived xenografts (PDXs) treated with PT2399 (data from ^26^ including sensitive PDXs only) for genes grouped by whether their expression in A498 cells is decreased by *shARNT* and not by *shBMAL1* (ARNTsp, yellow), by *shBMAL1* and not by *shARNT* (BMAL1sp, purple), by either *shARNT* or *shBMAL1* (Overlap, salmon), or neither (NA, gray). **** P < 0.0001 by two-way ANOVA with Tukey’s correction. Boxes depict the median and interquartile range (IQR), whiskers extend either to the minimum or maximum data point or 1.5*IQR beyond the box, whichever is shorter. Outliers (values beyond the whisker) are shown as dots. (**G-I**) Volcano plots depicting the expression changes for individual genes in the groups depicted in (*F*). Genes with padj < 0.05 are colored in red (fold change > 1.5) or blue (fold change < 0.67). (**J,K**) GOBP (*J*) or KEGG (*K*) pathways enriched among ARNT-specific or BMAL1-specific target genes in A498 cells. (**L,M**) Enrichment plots showing the impact of *shARNT* (*F*) or *shBMAL1* (*G*) on genes in the Hallmark HYPOXIA gene set.

**Figure S7:**
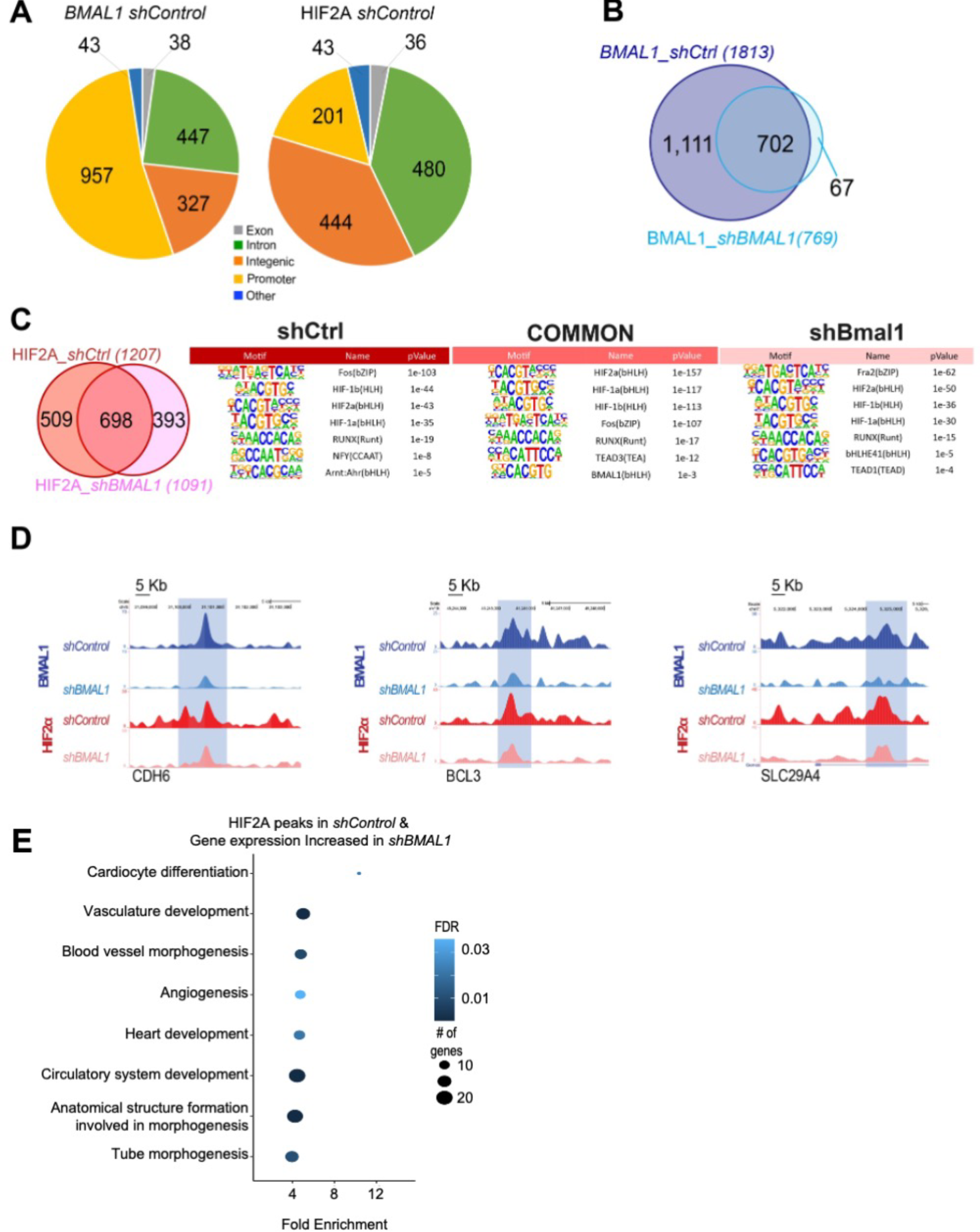
Impact of *shBMAL1* on HIF2α chromatin interactions. (**A**) Pie chart distribution of HOMER peak annotation for BMAL1 and HIF2α peaks in 786O cells expressing *shControl*. (**B**) Venn diagram depicting the numbers of genomic sites identified in CUT&RUN samples prepared using BMAL1-specific antibodies from 786O cells expressing *shControl* or *shBMAL1. (***C**) Left: Venn diagram depicting numbers of genomic sites identified in CUT&RUN samples prepared using HIF2α-specific antibodies from 786O cells expressing *shControl* or *shBMAL1*. Right: Top motifs enriched in chromatin associated with HIF2α in 786O cells expressing *shControl, shBMAL1*, or both (common) (D) Representative genome browser tracks for BMAL1 and HIF2α CUT&RUN signatures, showing *CDH6, BCL3,* and *SLC29A4* loci. (E) Data integration of RNA-seq and CUT&RUN HIF2α peaks. The x-axis represents the Enrichment Ratio, and the y-axis represents enriched GOBP pathways for genes associated with HIF2α peaks identified in CUT&RUN data from 786O cells expressing *shControl* with at least 5 genes, FDR < 0.05, and fold enrichment > 2. Expression of these genes was increased in 786O cells expressing *shBMAL1* compared to 786O cells expressing *shControl*. There were no significant KEGG pathway enrichments detected for genes with associated with HIF2α peaks identified in CUT&RUN data from 786O cells expressing *shControl* and no significant enrichment for either KEGG or GOBP databases for genes associated with HIF2α peaks identified in CUT&RUN data from 786O cells expressing *shBMAL1* with either increased or decreased expression in 786O cells expression in BMAL1-depleted 786O cells.

**Figure S8.**
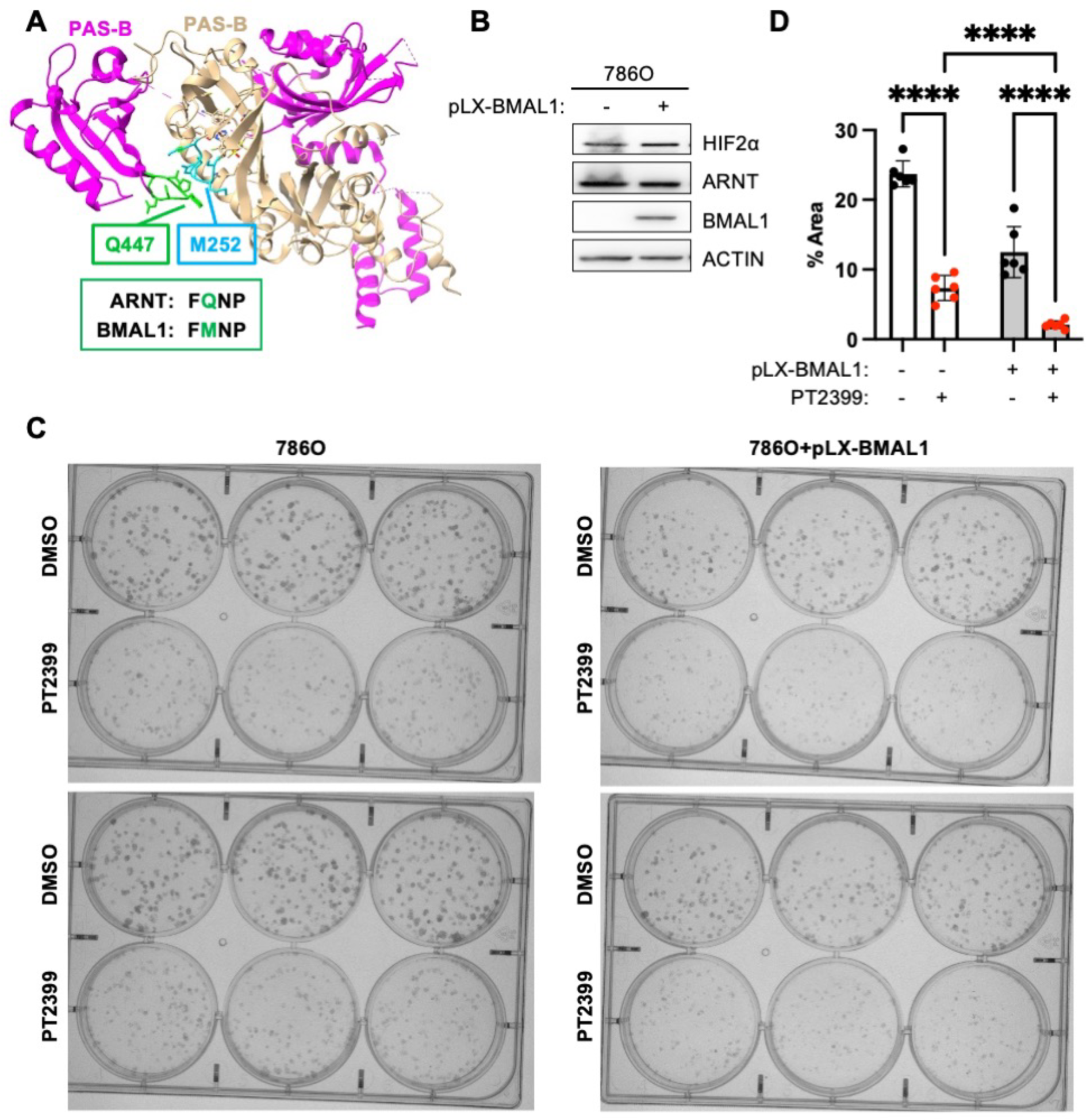
BMAL1-HIF2α heterodimers are sensitive to disruption by PT2399. (A) Three dimensional structure (PDB ID: 7W80) containing the bHLH and PAS domains of ARNT (magenta) and HIF2α (beige) bound to PT2399. Ligand binding to the PAS-B domain of HIF2α leads to structural reorganization of HIF2α loop (shown in blue), which thereby clashes with a loop in ARNT (shown in green). (B) Detection of HIF2α, ARNT, BMAL1, and ACTIN by immunoblot in 786O cells with (+) or without (-) lentivirus-mediated overexpression of BMAL1. (**C,D**) Representative images (*C*) and quantification (*D)* of colonies stained with crystal violet 10-16 days after plating 250 cells expressing the indicated plasmids per well in media containing vehicle (DMSO) or PT2399. Data in (*D*) represent the mean ± s.d. for six wells per condition from one experiment representative of at least three replicates. **** P < 0.0001 by two-way ANOVA with Tukey’s correction for multiple hypothesis testing.

## Notes

### Competing Interest Statement

The authors have declared no competing interest.

